# Offspring effect of maternal preconception exposure to metabolic disruptors

**DOI:** 10.1101/2025.05.22.655630

**Authors:** Carlos Diaz-Castillo, Stephanie R. Aguiar, Raquel Chamorro-Garcia

**Author notes:** Corresponding authors: Carlos Diaz-Castillo and Raquel Chamorro-Garcia.

## Abstract

Over the past several decades, there has been a resurgence in the field of multigenerational research, which focuses on the transmission of environmental effects across generations and their correlation with the prevalence of chronic diseases. Traditionally, it has been hypothesized that the propagation of such effects is mediated by alterations in gene regulatory elements sensitive to environmental cues. These alterations can be perpetuated throughout development and across generations in the absence of any genetic changes, such as DNA methylation or histone modifications or non-coding RNAs. Evidence suggesting that the compartmentalization of eukaryotic genomes into heterochromatin and euchromatin plays a crucial role in mediating multigenerational metabolism-disrupting effects elicited by the exposure to different metabolism disruptors has been observed in mice and fruit flies. This suggests that eukaryotic nuclear genomes may generally possess the capacity to integrate the impact of environmental cues in a metastable manner that is phenotypically relevant. In this study, we present the initial results of a murine model to determine whether preconception exposure to metabolism disruptors leads to metabolic alterations in the offspring of exposed individuals. We specifically focused on investigating the effects of three metabolism disruptors of distinct nature, designed to model the complexity of human exposures. Our findings are not only consistent with our central hypothesis but also open an unexpected avenue to explore whether preconception exposure to metabolism disruptors can predispose the offspring of exposed individuals to not only typical metabolic diseases such as obesity, but also to complex metabolic-psychiatric conditions such as anorexia.

## Introduction

Over the last 20 years, there has been increasing evidence showing how environment agents, such as chemical, biological, physical or psychosocial factors, play a critical role in the prevalence of chronic noncommunicable diseases in humans^1^. This shift in our understanding on the origin of disease, from heritable genetics to a gene-environment interaction approach, emerged after the recognition that only a small proportion of major chronic diseases, such as cancer and cardiovascular disease, can be attributed to genetic factors, <22% and <8%, respectively^2^. Animal model research has substantiated the significant impact of environmental exposures on health outcomes for exposed individuals and their subsequent generations^3^ .

Obesity is an example of chronic metabolic condition whose prevalence has dramatically increased in the last 50 years, and only ∼2.7% of cases can be attributed to genetic factors^4^. Exposure to metabolism disruptors, including pesticides, stress, and microorganisms, increases the susceptibility to developing chronic metabolic conditions, such as obesity, which can be propagated across multiple generations^5^. These exposures can alter lipid metabolism, energy balance, appetite regulation, and glucose homeostasis by interfering with hormone signaling pathways, including those involving insulin, leptin, and nuclear receptors^6^ .

Starting in 2013, we contributed to demonstrate that exposure to the biocide tributyltin (TBT) in female mice one week prior to conception and during the *in utero* development of their offspring, but not during fertilization to avoid exposing males, resulted in sexually dimorphic metabolic alterations in at least four offspring generations (F1, F2, F3, and F4)^7–10^. It is frequently debated whether the observed effects in the F1 and F2 offspring of females exposed to specific environmental factors during gestation could be directly attributable to these factors reaching the somatic tissues and germ cells of the F1 embryos developing *in utero*, respectively^11,12^. However, any effect observed in the F3 generation of descendants of exposed females and beyond could not have been directly triggered by the environmental factor in question but must have been mediated by perturbations that can be propagated through development and across generations^11,12^. To elucidate the underlying mechanism mediating the transgenerational metabolic disruption induced by TBT, we devised a series of integrative analyses of several traits related to the epigenomic function^8,9^. Specifically, we compared epigenetic marks such as DNA methylation at a genomic scale and other structural and genomic traits known to be regulated by epigenetic modifications, including chromatin accessibility and gene expression, in the somatic tissues and mature male gametes of the F3 and F4 descendants of females that had been exposed to TBT and controls^8,9^. In most of our analyses, we detected a dichotomy for TBT-dependent alterations with regard to the heterochromatic and euchromatic compartments of the mouse genome; changes in a particular trait were significantly biased in one direction in heterochromatin and the opposite direction in euchromatin^8,9^. In conjunction with our investigation of the distribution within the mouse genome of genes regulating metabolism and chromatin organization, we proposed that the most plausible mechanism underlying the TBT-dependent multigenerational effects is that TBT triggered a perturbation of chromatin organization with the ability to self-reconstruct through development and across generations and predispose the progeny of exposed females to metabolic disorders^8,9^. Although our analyses did not directly address which layers of chromatin organization were altered upon TBT exposure, our findings were consistent with the possibility that such perturbation encompassed an alteration of the heterochromatin/euchromatin compartmentalization (HEC)^8,9^.

To the best of our knowledge, the ability of HEC to function as an epigenetic mechanism of genome regulation capable of mediating multigenerational metabolic disruptions has only been demonstrated in a single study on *Drosophila melanogaster*. In 2014, Öst et al. observed a similar dichotomy in gene expression alterations for heterochromatic and euchromatic genes in the offspring of male *Drosophila melanogaster* subjected to diets with varying sugar content^13^. Exposure to these diets also led to alterations in metabolic traits and the expression of heterochromatin markers^13^. Notably, the association between the dichotomy of perturbations in epigenomic function traits and metabolic disorders in the descendants of individuals exposed to environmental modulators of metabolism in two independent studies that otherwise differ in the nature, duration, and sex of exposure, the number of generations separating exposed and descendants, and even the epigenetic characteristics of the species under study, *Drosophila melanogaster,* has very low levels of DNA methylation^14^, could suggest that eukaryotic HEC is generally susceptible to modulation by environmental exposure to metabolism disruptors in a metastable manner, resulting in metabolic effects in the descendants of exposed individuals^8,9,13^.

Based on existing knowledge regarding the developmental establishment of HEC and the relevance of human preconception exposures eliciting multigenerational chronic metabolic diseases, we hypothesized that preconception exposure to metabolism disruptors can predispose to complex metabolic diseases in the offspring of exposed individuals through HEC perturbations.

On the one hand, research conducted across multiple species employing diverse methodologies has demonstrated that the earliest indications of genome compartmentalization and heterochromatin formation emerge very early in metazoan development, predating the full activation of the zygotic genome that characterizes the maternal-to-zygotic transition (MTZ)^15,16^. Furthermore, the study of reporters of heterochromatic function in *Drosophila melanogaster* has suggested that adult heterochromatin structures are established very early in development and are faithfully propagated throughout development^17^. Recent theoretical modeling for the propagation of heterochromatin-related epigenetic marks indicates that the spatial compartmentalization of heterochromatin and the limited availability of enzymes essential for its maintenance are necessary for the faithful propagation through cell differentiation and development of heterochromatin-related structures^18^. Given that during the MTZ transition, the zygote is largely transcriptionally silent^19^, all processes occurring along MTZ, including the establishment of heterochromatin, occur at the expense of the limited amounts of material deposited in the oocyte and to a lesser extent in the sperm. Considering these factors, it appears plausible that any environmental exposure resulting in alterations of the gamete material required for the establishment of the zygotic heterochromatin subsequently led to perturbations of heterochromatin that are propagated throughout development, thereby mediating phenotypes that reflect the biased localization of genes with specific functionalities in relation to heterochromatin^9,13^.

On the other hand, although human multigenerational studies are limited, the few that exist support the hypothesis that exposure to environmental stressors before conception contributes to chronic conditions in their offspring of at least two generations The Dutch Famine stands as a pivotal human study that elucidates the significance of identifying critical windows of susceptibility to environmental stressors and chronic disease prevalence in subsequent generations^20^. Numerous analyses of this cohort have demonstrated that low calorie intake during early, mid, and late gestation can result in chronic metabolic, psychological, and immunological disorders^20^. Notably, individuals exposed to famine between 10 weeks before and 10 weeks after conception (*i.e.,* preconception and early gestation exposure) exhibited a substantial increase in obesity, coronary heart disease, and elevated blood cholesterol levels compared to those exposed to famine during mid and late gestation^20^. Other human studies have shown that alterations in food availability or tobacco use during puberty have an effect in their grand offspring^21,22^. Altogether, these findings underscore the profound impact of preconception and early gestation events on transgenerational health and disease outcomes.

We here present the results of our initial study investigating whether preconception exposure of female mice to three diverse metabolism disruptors leads to metabolism perturbation in their offspring and address the susceptibility of HEC to accommodate such exposures in a metastable manner. We selected three metabolism disruptors, TBT, inorganic arsenic (iAs) and diet, to model real-life human exposures of very different nature. TBT is an example of a human-made metabolism-disrupting chemical commonly found in soil, dust, and water and in human samples from liver, placenta, and blood, that has been associated with multigenerational obesity^7–10,23^. iAs is a naturally-occurring element ubiquitously found in soils, sediment, and groundwater, and it poses major threats to global public health as around 200 million people worldwide is exposed to high concentrations^24^. Prenatal exposure to arsenic has been associated with increased adiposity and increased risk of type-2 diabetes in human populations^25^. Maternal preconception exposure to environmentally relevant doses of iAs in mice led to transgenerational metabolic perturbations, but little is known about potential epigenetic alterations associated with such transgenerational phenotypes^26^. Lastly total Western diet (TWD), whose micro- and macro-nutrient content represents the diet 50% of the U.S. population take, is an example of life-style metabolism disruptor^27^.

It is imperative to recognize that humans are subjected to a multitude of environmental stressors of varying nature from the moment of conception until death. This concept is encapsulated within the exposome framework^28^, which encompasses the intricate interplay between life experiences and exposures that contribute to modulating susceptibility to disease. In this study, our primary objective was to elucidate the significance of preconception exposure to environmental stressors and their metabolic effect in the next generation. Although these exposures will not directly address the complexity of the exposome, it can be a stepping-stone in our understating of whether environmental factors humans are exposed to and that have been associated with metabolic disorders, do so via similar mechanisms of action.

## Methods

### Mouse work

We conducted all mouse procedures at the University of California, Santa Cruz (UCSC) Vivarium. These procedures were reviewed and approved by the UCSC Institutional Animal Care and Use Committee (UCSC IACUC) as part of the animal protocol Chamr1908.

We purchased 160 female and 80 male C57BL/6J mice from Jackson Laboratory (strain #000664). To ensure the acclimation of mice to the UCSC Vivarium, we scheduled their arrival one week prior to the beginning of the experiment. Upon arrival, we ear-punched and weighed female mice and subsequently housed them in four-mouse cages. To minimize any initial body weight disparities between the experimental groups, we randomly assigned mouse cages to five distinct groups. Initially, we ranked the mouse cages based on the cumulative body weights of the mice within each cage, from highest to lowest. Subsequently, we divided the ranked list into eight subgroups of five cages each and randomly assigned each of the five cages within each subgroup to one of the experimental groups.

We exposed 5-week-old female mice in each group to the following combinations of treated drinking water and diet for 3.5 weeks:

- DMSO (negative control) group: 0.1% Dimethylsulfoxide (DMSO) in water and control diet (CD; Envigo Teklad Diets TD.140148).
- 5TBT group: 5 nM tributyltin (TBT) and 0.1% DMSO in water and CD.
- 50TBT group: 50 nM tributyltin and 0.1% DMSO in water and CD.
- IAS group: 10 µg/L sodium meta(arsenite) (inorganic arsenic, iAs) and 0.1% DMSO in water and CD.
- TWD group: 0.1% DMSO in water and Total Western Diet (TWD; Envigo Teklad Diets TD.110919).

The Agency for Toxic Substances and Disease Registry (ATSDR) determined that the no-observed adverse effect level (NOAEL) for TBT is 0.025 mg/Kg/day^29^. 50 nM TBT in drinking water is equivalent to 0.005 mg/Kg/day assuming an average mouse weight of 30 g and an average of 10 mL of water taken per day. Thus, 5 and 50 nM TBT represents 50 and 5 times lower than the NOAEL, respectively. We used 10 µg/L of inorganic arsenic int eh drinking water as that is the allowable level established by the U.S. Environmental Protection Agency^30^. Total Western diet (TWD) represents the diet 50% of U.S. eat on a regular basis^27^.

We replenished the mouse feed weekly and water bottles semiweekly to maintain freshness and control for food and water consumption. Prior to initiating the exposure and subsequent weekly treatments, we weighed the mice before and after fasting for a period of four hours. To safeguard the reproductive health of female mice from the adverse effects of fasting, we did not fast them at the end of the exposure period and immediately before mating.

After 3.5 weeks of exposure at 8 weeks of age, we mated exposed female mice with unexposed same-age male mice for one week to generate their F1 offspring. We checked females daily and determined successful matings by observing copulatory plugs. When we identified the copulatory plug, mice were returned to four-mouse cages with their same cage mates from the exposure period for the rest of the pregnancy. Two days prior to the anticipated birth, we relocated pregnant females to individual cages equipped with ample bedding material, enabling them to construct a nurturing nest. We monitored closely births and the welfare of litters using the least invasive methods possible until they reached a sufficient age for further manipulations.

At 8-11 days of age, we toeclipped all F1 mice. At 3 weeks of age, we weaned F1 mice and selected at least 10 mice per sex and exposure group. We housed mice of the same sex and exposure group together. We weighed F1 mice at weaning and continued to do so weekly thereafter until the conclusion of the experiment. We fed F1 mice CD between 3 and 7 weeks of age and TWD between 7 weeks of age and the end of the experiment. Prior to changing diets at 7 weeks of age, we weighed F1 mice before and after fasting them for 12 hours. We measured fasting glucose using a Contour® blood glucose meter (BAYER) and Contour® blood glucose strips (BAYER) with a drop of blood drawn from the mouse tails by puncturing them with a needle. At 12 weeks of age, we fasted F1 mice for 12 hours and euthanized them using an overdose of isoflurane followed by cardiac exsanguination. We determined fasting glucose levels from peripheral blood as previously indicated and harvested gonadal and inguinal white adipose tissue (gWAT and iWAT, respectively) depots and liver. We draw blood directly from the heart using EDTA-flushed syringes to minimize coagulation and collected cardiac blood samples in microcentrifuge tubes with 6 µL of 100X for a protease inhibitors cocktail (Sigma Aldrich NC2042678). To separate plasma, we centrifuged blood samples at 5,000 x g for 10 minutes at 4°C. We weighed gWAT, iWAT, and liver samples and snap-frozen them in dry ice. We maintained plasma, gWAT, iWAT and liver samples frozen until required for downstream analyses. Dissections were performed by all available appropriately trained members of the Chamorro-Garcia group. To minimize the daily dissection workload, we distributed F1 mice into five groups, ensuring equal representation of each exposure group and sex. Each group underwent dissection over a period of five consecutive days. The lead experimenter (Diaz-Castillo) employed a randomized approach to assign the order of mouse dissection and assign each mouse to a dissector without prior knowledge of their group or sex.

### Metabolic analyses

We determined body weight by carefully placing each mouse in an empty beaker atop a standard laboratory balance. To minimize the confounding effect of age-related differences, particularly in younger F1 mice, we corrected body weight data by the days of age of each mouse. We assessed the significance of body weight differences between exposure groups (5TBT, 50TBT, IAS, and TWD) and the control group (DMSO) using unmatched-measures Monte Carlo-Wilcoxon (uMCW) tests (see Statistics section). Similarly, we assessed the significance of body weight changes associated with fasting challenges between exposure groups and controls using matched-bivariate Monte Carlo-Wilcoxon (mbMCW) tests (see Statistics section).

To determine water consumption, we filled water bottles in each cage with 200 mL of treated water (entry water) and measured the volume of unconsumed water after three or four days (exit water). Similarly, we weighed the amount of fresh food provided in each cage (entry food) and the unconsumed food after one week (exit food). We assessed the significance of entry and exit food and water differences between exposure groups and controls using mbMCW tests.

We corrected plasma glucose levels determined in fasted mice by the body weight and the days of age of each mouse. We assessed the significance of fasting glucose differences between exposure groups and controls using uMCW tests.

We weighed gWAT, iWAT and liver samples harvested from F1 mice using a precision balance. We corrected tissue weights by the body weight and the days of age of each mouse. We assessed the significance of tissue weight differences between exposure groups and controls using uMCW tests.

We submitted 50 µL of F1 plasma samples obtained during their euthanasia to Eve Technologies (Calgary, Canada) for the analysis of 11 metabolites included in the Mouse, Rat Metabolic Array (MRDMET), namely amylin, gastric inhibitory peptide (GIP), ghrelin, glucagon-like peptide 1 (GLP-1), insulin, leptin, peptide YY (PYY), glucagon, pancreatic peptide (PP), resistin, and connecting perptide (C-Peptide). We set to zero any data outside of the range of the standard curve or that could not be extrapolated mathematically. We corrected plasma levels of each metabolite by the body weight and the days of age of each mouse. We assessed the significance of differences in plasma levels among exposure groups and controls using uMCW tests. We performed Principal Component Analysis (PCA) to determine the similarities between mouse groups by utilizing data for metabolites that were determined for all female or male mice. We used the function *prcomp* from the R package *Stats* (version 4.5)^31^ to perform the PCA analyses, and the functions *fviz_eig* and *fviz_pca* from R package *Factoextra* (version 1.0)^32^ to visualize the results.

### Transcriptomics

We used a VWR® Micro Homogenizer (catalog number #10032-328) and Qiagen RNeasy Plus kits (catalog number #74034) to extract RNA from F1 gWAT and liver, following the manufacturer’s protocols. Subsequently, we submitted the RNA samples to the University of California, Irvine (UCI) Genomics High Throughput Facility for the preparation of sequencing libraries using Illumina TruSeq Stranded mRNA kits, and the sequencing of these libraries using an Illumina NovaSeq 6000 sequencer. Libraries were sequenced up to three times to obtain approximately 30 million 150 bp paired reads per sample. In total, 100 RNA samples were sequenced (2 sexes x 2 tissues x 5 conditions x 5 replicates per condition).

To proceed with the analysis of the sequencing results obtained from UCI Genomics High Throughput Facility we used the following accessory files and three sets of informatics tools.

#### Accessory files for RNA-seq analyses

We downloaded fasta files containing the sequences of all major autosomes and the *X* chromosome, and the file “mm39.ncbiRefSeq.gtf.gz” which includes the primary gene transcript annotation for the June 2020 GRCm39/mm39 assembly of the mouse genome from the UCSC Genome Browser (https://hgdownload.soe.ucsc.edu/downloads.html#mouse)^33^. Furthermore, we retrieved the most recent core Gene Ontology from the Gene Ontology website (https://geneontology.org/)^34,35^.

#### Linux/Python tools

We used the *isoSegmenter* tool (https://github.com/bunop/isoSegmenter)^36^ to define isochores and isochore classes for the mouse genome. We executed *isoSegmenter* with the fasta files corresponding to individual chromosomes, which we had downloaded from the UCSC Genome Browser. In essence, *isoSegmenter* operates in three phases. Initially, it segments the sequence for each chromosome into non-overlapping 100 kb windows. Subsequently, it assigns these windows to five classes based on their GC content. Lastly, it defines isochores by concatenating juxtaposed windows belonging to the same class. The five isochore classes exhibit the following GC contents: L1 isochores have a GC content below 37%, L2 isochores have a GC content between 37% and 41%, H1 isochores have a GC content between 41% and 46%, H2 isochores have a GC content between 46% and 53%, and H3 isochores have a GC content exceeding 53%.

#### Galaxy Platform tools

We uploaded the fastq files obtained from the UCI Genomics High Throughput Facility to the Galaxy Platform website (usegalaxy.org)^37^. We used the Galaxy Platform tools *FastQC* (version 0.12) and *multiQC* (version 1.27)^38^ to assess the quality of reads for each sample and generated multi-sample reports, respectively. We used the Galaxy Platform tool *Cutadapt* (version 5.0)^39^ to remove adapter sequences. We used the Galaxy Platform tools RNA Star (version 2.7)^40^ and featureCounts (version 2.0)^41^ to map and count reads to mouse genes, respectively, employing the annotation file “mm39.ncbiRefSeq.gtf.gz” to map reads to splice junctions. We used the Galaxy Platform tool Intersect (version 1.0) to determine the overlap of genes and isochores in the mouse genome. Upon the completion of all Galaxy Platform operations, we downloaded a table including RNA-seq gene-wise read counts for each gWAT and liver sample and isochore class associations.

#### R tools

We conducted all transcriptomic analyses using R (version 4.5)^31^ packages in Rstudio (Version 2024.12.1+563)^42^. We used base functions of R and the package *data.table* (version 1.17)^43^ to perform all data operations unless stated otherwise.

We used the R package *ComplexHeatmap* (version 2.24)^44^ to visualize the expression patterns across samples of the 100,000 more expressed genes. We aggregated all raw counts for each gene across all samples and identify genes with the top 100,000 cumulative counts. Subsequently, we normalized gene read counts to counts per million (cpm) by dividing raw read counts for each gene by the sum of raw read counts for all genes in each sample and multiplying the result by 1 million. To rescale normalized counts for each gene across all samples to the same 0-1 range, we employed the following formula: (*x* - min(*x*)) / (max(*x*) - min(*x*)), where *x* represents cpm values for each gene and sample, and min(*x*) and max(*x*) denote the minimum and maximum cpm values for each gene across samples, respectively.

To assess the significance of gene expression differences between exposure groups and controls, we performed uMCW tests (see Statistics section). We extracted read count tables for each of the 16 exposure group *versus* control contrast combinations (2 sexes x 2 tissues x 4 exposure groups). We considered a gene to be expressed if at least 2 sequencing reads had been mapped to this gene for at least 2 samples in each exposure group *versus* control contrast and discarded non-expressed genes. We normalized gene read counts to cpm as previously indicated.

To determine whether genes with the most extreme differences in expression between exposure groups and controls are significantly associated with specific functionalities, we performed pre-ranked Gene Set Enrichment Analyses (GSEA)^45^ using the R package *fgsea* (version 1.34)^46^ and the most updated mouse Gene Ontology (GO) annotation obtained from R package *msigdbr* (version 10.0)^47^. We restricted our *fgsea* analyses to GO sets of the Biological Process (BP) category with at least 15 and less than 500 genes. For each exposure group *versus* control contrast, we ranked genes using the uMCW bias indexes (BIs) from the highest to the lowest value. Thus, the top of the list will be populated by genes that are more expressed in the exposure group than in controls, and the bottom of the list will be populated by genes that are more expressed in controls than in the exposure groups. GSEA calculates an enrichment score (ES) representing the extent to which any GO-BP set is overrepresented at either extreme of a pre-ranked list of genes, a normalized enrichment score (NES) to account for the size of the set, and a *P* value calculated by comparing observed ES with a null distribution of ES obtained after repeatedly permuting gene ranks^45^.

To assess the significance of concerted changes in expression for genes in the whole transcriptome and specific regions of the genome such as chromosomes, individual isochores and isochores of the same class, we performed biased-measures Monte Carlo-Wilcoxon (bMCW) tests (see Statistics section). To restrict these analyses to genes encoded in the main chromosomes of the nucleus for both females and males, we filtered out genes located in the mitochondrial genome, the *Y* chromosome and unassembled chromosome segments from the table including RNA-seq gene-wise read counts for all samples and isochore class associations we downloaded from the Galaxy Platform. We further filtered out non-expressed genes and normalized read counts for each sample as previously indicated.

We determined the relative fraction of genes overlapping isochores of each class for genes in each autosome and the *X* chromosome and for genes associated to specific GO-BP terms for each exposure group *versus* control contrast. For each chromosome and GO-BP term and each isochore class, we calculated relative gene fractions as log_10_((*x*/*n*) / (*X*/*N*)), where *x* and *X* represent the number of genes in each chromosome or associated with each GO-BP term overlapping isochores of each class or any class, respectively, and *n* and *N* represent the number of genes for the whole transcriptome overlapping isochores of each class or any class, respectively.

We used the R package *smplot2* (version 0.2)^48^ to visualize the outcomes of Pearson correlation analyses between leptin plasma levels and *Lep* gene expression in gWAT, as well as between the expression of *Lep* and *Lnc-Lep* genes in gWAT.

### Other informatic tools

We used the following R packages to process data, visualize results, and prepare figures for publication: *data.table* (version 1.17)^43^, *ggplot2* (version 3.5)^49^, *ggplotify* (version 0.1)^50^, *ggrepel* (version 0.9)^51^, *ggtext* (version 0.1)^52^, *patchwork* (version 1.3)^53^, *RColorBrewer* (version 1.1)^54^, *scales* (version 1.3)^55^, and *svglite* (version 2.1)^56^. Upon peer-reviewed publication, we will include copies of any informatics codes needed to reproduce our analyses and figures.

### Statistics

To ascertain the statistical significance of differences observed between exposure groups and controls for litter, metabolic, and transcriptomic traits in the most comprehensive manner, we conducted most of our analyses using Monte Carlo-Wilcoxon (MCW) tests. Initially, MCW tests were devised to ascertain whether a set of matched-paired quantitative measures exhibited a statistically significant bias in the same direction when compared with what would be expected by chance^8,9,57–59^. Recently, we have reformulated MCW tests to interrogate four distinct data structures and developed the R package *MCWtests*^60^ to publicly distribute the functions that facilitate conducting such tests. A comprehensive description of MCW testing rationale and specific MCW tests can be found at https://diazcastillo.github.io/MCWtests/index.html. Briefly, all MCW tests follow two basic steps. Firstly, to quantitatively determine the magnitude and direction of the bias between two conditions for the measure being analyzed, MCW tests compute a bias index (BI) that spans the range of 1 to -1, indicating that the measure is completely biased in each conceivable direction. Subsequently, to ascertain the statistical significance of the BIs computed for the user-provided dataset (observed BIs), a series of expected-by-chance BIs are generated by repeatedly rearranging the original dataset and computing BIs for each iteration. *P_upper_* and *P_lower_* values are subsequently calculated as the proportions of expected-by-chance BIs that exhibit values equal to or higher than and equal to or less than the observed BIs, respectively.

MCW tests are particularly suitable for highly integrative studies such as this one. On the one hand, like other non-parametric approaches, MCW tests do not necessitate the original data to adhere to any specific distribution or require any mathematical transformation to conform the data to a predefined distribution. On the other hand, since the bias information (BI) values always range between -1 and 1, comparing results from MCW tests conducted with data from different ranges, scales, or even traits become straightforward.

In our study, we employed three of the four MCW tests provided by the R package MCWtests: unmatched-measures MCW (uMCW) tests, matched-measures bivariate MCW (mbMCW), and biased-measures MCW (bMCW) tests.

uMCW tests assess whether two sets (*e.g.*, *a* and *b*) of unmatched measures exhibit significant bias in the same direction. The uMCW testing process involves the following steps. First, all possible disjoint data pairs using measures from both sets are drawn. Second, the second measure in each pair is subtracted from the first measure. Third, measure pairs are ranked based on the absolute value of their differences from lowest to highest. Measure pairs whose subtraction equals 0 are assigned a 0 rank. Measure pairs with the same difference values are assigned the lowest rank. Fourth, ranks are signed using the sign of the measure differences calculated in the third step. Fifth, signed ranks for each type of set subtractions (*e.g.*, *a-b* and *b-a*) are aggregated. Sixth, bias indexes (BIs) are calculated as the sum of signed ranks divided by the maximum value this sum would have if all measures in the first set (*e.g.*, *a*) were greater than those in the second set (*e.g.*, *b*). Finally, the significance of observed BIs is determined by comparing them with a collection of expected-by-chance BIs calculated after repeatedly rearranging the measure assortment between the two sets, and calculating *P_upper_* and *P_lower_* values as previously indicated. uMCW tests can follow two paths based on the user-defined parameter *max_rearrangements.* If the number of potentially distinctive data rearrangements is less than *max_rearrangements*, uMCW tests will draw all potentially distinctive rearrangements, and *P_upper_* and *P_lower_* values will represent an exact estimation of the significance of observed BIs. If the number of potentially distinctive data rearrangements is greater than *max_rearrangements*, uMCW tests will perform a number of rearrangements equal to *max_rearrangements*, and *P_upper_* and *P_lower_* values will represent an approximated estimation of the significance of observed BIs.

mbMCW tests assess whether two sets (*e.g.*, *x* and *y*) of inherently matched-paired measures are significantly differentially biased in the same direction. The mbMCW testing process involves the following steps. First, for each matched-pair of measures, the two possible subtractions are calculated (*e.g.*, *a-b* and *b-a*). Second, for each type of subtractions (*e.g.*, *a-b* and *b-a*), measure pairs are ranked from lowest to highest. Measure pairs whose subtraction equals 0 are assigned a 0 rank. Measure pairs with the same difference values are assigned the lowest rank. Third, ranks are signed using the sign of the measure differences calculated in the first step. Fourth, signed ranks for each type of subtractions (*e.g.*, *a-b* and *b-a*) and sets of paired measures (*e.g.*, *x* and *y*) are aggregated. Fifth, bias indexes (BIs) are calculated as the sum of signed ranks divided by the maximum value that sum would have if all measure differences in the first set (*e.g.*, *x*) were greater than those in the second set (*e.g.*, *y*). Finally, the significance of observed BIs is determined by comparing them with a collection of expected-by-chance BIs calculated after repeatedly rearranging of the measure matched-pair assortment between the two sets, and calculating *P_upper_* and *P_lower_* values as previously indicated. mbMCW tests can also follow two paths to calculate exact and approximated estimations of the significance of observed BIs as indicated for uMCW tests.

bMCW tests are a combination of two tests that assess whether a set of measures of bias for a quantitative trait between two conditions and a specific subset of these bias measures are significantly biased in the same direction. The bMCW testing process involves the following steps. First, all bias measures are ranked using their absolute values from lowest to highest. Second, ranks are signed using the sign of the bias measure under study. Third, a whole-set bias index (wBI) is calculated by summing signed ranks for all elements in the dataset and dividing it by the maximum number this sum could have if all bias measures were positive. Third, for each subset of elements in the dataset under study, a subset bias index (sBI) is calculated by summing signed ranks for the elements in the subset and dividing it by the maximum value this sum would have if the elements in the subset had the most positive values in the whole dataset. Fourth, the significance of observed wBIs is determined by comparing them with a collection of expected-by-chance wBIs calculated after repeatedly rearranging the signs of all signed ranks, and calculating *P_upper_* and *P_lower_* values as previously indicated. Finally, the significance of observed sBIs is determined by comparing them with a collection of expected-by-chance sBIs calculated after repeatedly rearranging the number of elements in the whole dataset that are associated with the subset under study, and calculating *P_upper_* and *P_lower_* values as previously indicated. bMCW tests can also follow two paths to calculate exact and approximated estimations of the significance of observed wBIs and sBIs as indicated for uMCW tests.

We used uMCW tests to determine whether mouse body weights, fasting glucose, tissue weights, plasma metabolite levels, litter size and sex ratio, and gene expression for individual genes were significantly biased in the same direction for each exposure group when compared with controls. We used mbMCW tests to determine the significance of fasting body weights and water and food consumption for each exposure group when compared with controls. In the case of fasting body weights, we conducted mbMCW tests for mouse pre-fasting and post-fasting body weights data pairs for each experimental group. In the case of water and food consumption, we conducted mbMCW tests for mouse cage mouse cage entry and exit, water and food data pairs for each experimental group. We used bMCW tests to determine whether gene expression differences between each exposure group and controls for genes in the whole transcriptome, overlapping specific isochores, overlapping isochores of the same class or located within autosomes and the *X* chromosome were significantly biased in the same direction. The measure for gene expression difference we used for bMCW tests was the uMCW test BIs obtained for each gene and exposure group *versus* control contrast. For all our MCW tests, we set *max_rearrangements* parameter to 10,000.

In this study, we also used other statistical analyses, including principal component analysis (PCA), gene set enrichment analysis (GSEA), and Pearson correlations, as previously described in the Metabolic and Transcriptomics section.

## Results

### Direct exposure to TBT, inorganic arsenic and total western diet cause different metabolic disruptions

We randomly divided 160 female C57BL/6J mice into five equal-sized groups. Each group was exposed to one of the following combinations of treated drinking water and diet for 3.5 weeks, starting at 5 weeks of age: DMSO and CD (negative control), 5 nM TBT and CD (5TBT group), 50 nM TBT and CD (50TBT group), 10 µg/L sodium (meta)arsenite and CD (IAS group), and DMSO and TWD (TWD group). To assess the efficacy of these exposures and determine if 5TBT, 50TBT, IAS, and TWD exposures caused metabolic disruptions compared to DMSO controls without excessively disturbing the exposed mice, we repeatedly measured body weight, fasting body weight, water, and food consumption. We weighed F0 females before and after a 4- hour fasting challenge immediately before starting the exposures (timepoint T0) and weekly thereafter for three weeks (Figure 1A). We determined water consumption twice a week after the start of the exposure by measuring the volume of entry water deposited in each cage glass bottle in the preceding timepoint and the volume of exit water left unconsumed in each bottle since then (Figure 1A). We determined food consumption weekly after the start of the exposure and at week 3.5 of the exposure period by measuring the weight and caloric value of entry food deposited in each cage in the preceding timepoint and the weight and caloric value of exit food left unconsumed in each since then (Figure 1A). We weighed female mice immediately prior to mating but did not subject them to fasting challenges to avoid exposing them to a stress that could jeopardize their fertility (Figure 1A).

**Figure 1.**
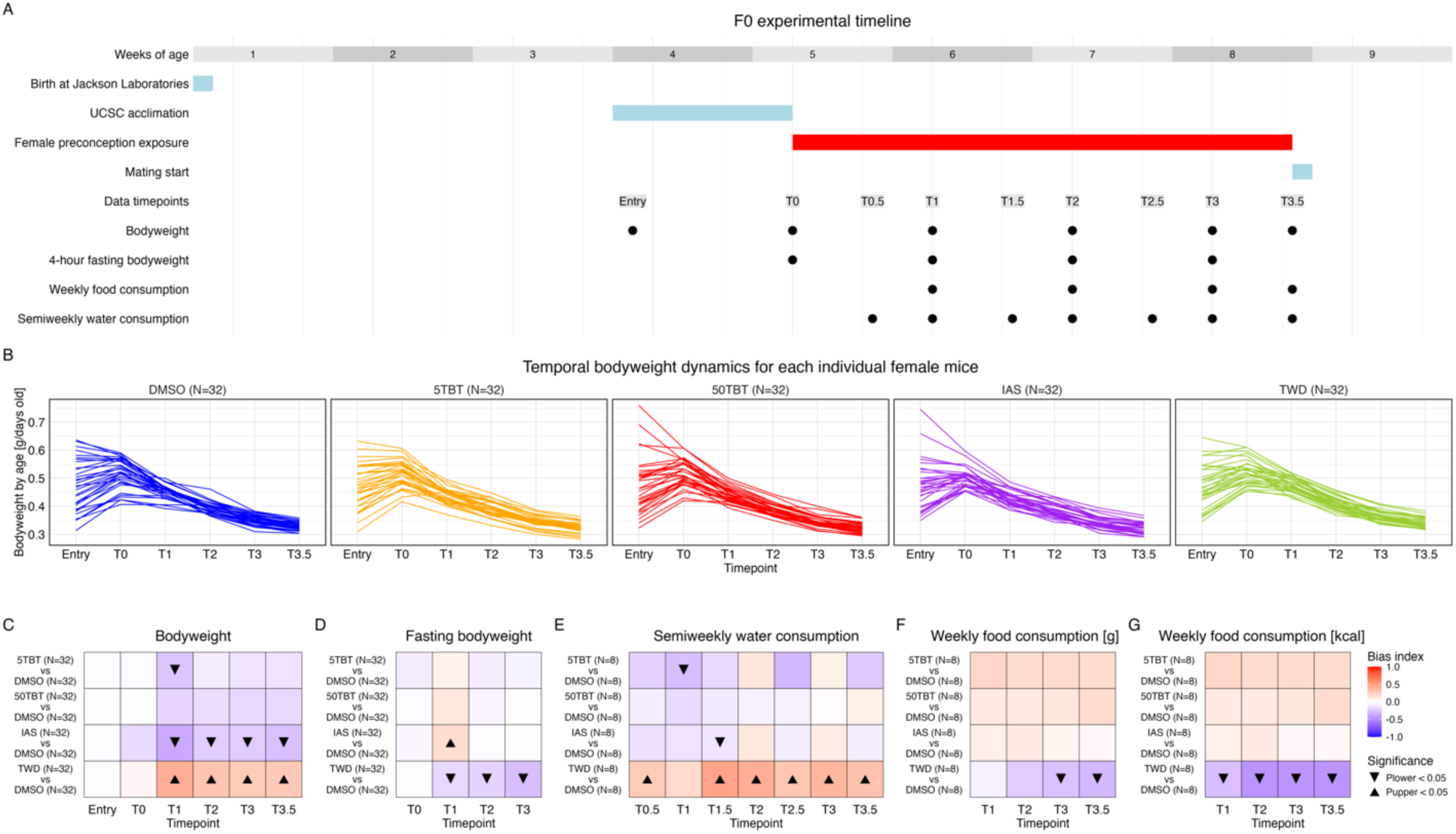
Direct effect of the exposure of F0 female mice to TBT, inorganic arsenic, and Total Western Diet. **A.** Experimental timeline detailing primary mouse operations involving exposure to TBT, iAs, and TWD in female mice, along with the timepoint structure employed to evaluate variations in body weight, fasting body weight, water, and food consumption between exposure groups and controls. **B.** Body weight dynamics for each mouse within each experimental group, spanning their entry at UCSC Vivarium to the conclusion of the exposure period at timepoint T3.5. **C.** Comparison of body weight normalized by age (g/days old) between exposure groups and controls using uMCW tests. **D.** Comparison of fasting body weight normalized by age (g/days old) between exposure groups and controls using mbMCW tests. **E.** Comparison of water consumption (mL) between exposure groups and controls using mbMCW tests. **F.** Comparison of food mass consumption (g) between exposure groups and controls using mbMCW tests. **G.** Comparison of food caloric consumption (kcal) between exposure groups and controls using mbMCW tests. The number of replicates (N) for each experimental group is indicated in each panel.

To determine whether body weights, fasting body weights, water consumption and food consumption were significantly different between 5TBT, 50TBT, IAS and TWD groups when compared with the control DMSO group, we used Monte Carlo-Wilcoxon (MCW) tests (see Methods for more details). Briefly, to quantitatively determine the magnitude and direction of the bias between two conditions for the measure being analyzed, MCW tests compute a bias index (BI) that spans the range of 1 to -1, indicating that the measure is completely biased in each conceivable direction. Subsequently, to ascertain the statistical significance of the BIs computed for the user-provided dataset (observed BIs), a series of expected-by-chance BIs are generated by repeatedly rearranging the original dataset and computing BIs for each iteration. *P_upper_* and *P_lower_* values are subsequently calculated as the proportions of expected-by-chance BIs that exhibit values equal to or higher than and equal to or less than the observed BIs, respectively. MCW tests include four different tests to interrogate different data structures. Unmatched- measures MCW (uMCW) tests assess whether two sets (*e.g.*, *a* and *b*) of unmatched measures exhibit significant bias in the same direction. We used uMCW tests to determine whether mouse body weights were significantly biased in the same direction for each exposure group when compared with controls. Matched-measures bivariate MCW (mbMCW) tests assess whether two sets (*e.g.*, *x* and *y*) of inherently matched-paired measures are significantly differentially biased in the same direction. We used mbMCW tests to determine the significance of fasting body weights and water and food consumption for each exposure group when compared with controls. In the case of fasting body weights, we conducted mbMCW tests for mouse pre-fasting and post-fasting body weights data pairs for each experimental group. In the case of water and food consumption, we conducted mbMCW tests for mouse cage mouse cage entry and exit, water and food data pairs for each experimental group.

Although 5TBT, 50TBT, IAS, and TWD exposures appeared to disrupt the metabolism of exposed females, these disruptions manifested differently depending on the exposure. 5TBT exposure resulted in a significant reduction in body weight immediately after exposure, which was subsequently mitigated (Figure 1C). 50TBT exposure also resulted in a reduction in body weight, but such a difference was not significant at any timepoint. No consistent patterns were observed for fasting body weight or water and food consumption (Figure 1D-G) for either TBT exposure. While TBT mice exhibited a tendency to consume more food and drink less water compared to DMSO controls, water consumption was only significantly decreased for the 5TBT group in a single timepoint (Figure 1E-G).

IAS exposure also led to a substantial reduction in body weight immediately after exposure, but this effect persisted throughout the entire exposure period (Figure 1C). Additionally, during the initial week of exposure, but not thereafter, fasting body weight in IAS females was significantly higher compared to DMSO controls (Figure 1D). This observation suggests that the reduction in body weight observed in IAS females may not have been equally responsive to the fasting challenge as the controls during the time when the body weight differences were most pronounced. Furthermore, IAS females appeared to consume more food and drink less than controls throughout the exposure period, although these differences were rarely significant (Figure 1E-G).

The group exhibiting the most pronounced and substantial differences was the TWD group. These female mice demonstrated significantly higher body weights, lower fasting body weights, increased water consumption, and decreased food consumption compared to controls throughout the exposure period (Figure 1C-G). The body weight excess in TWD mice is more readily mobilized even during a mild fasting regimen, which aligns with the substantial increase in water consumption observed in TWD mice when compared to controls.

All these patterns are consistent with TBT and IAS exposures causing similar decreases in body weight, with the IAS group weight loss being more pronounced, and TWD exposure causing an increase in body weight. The fact that these four exposures lead to distinct metabolic disruptions provides a great opportunity to comparatively assess if they lead to distinctive metabolic disruptions in the offspring of exposed females.

### Preconception exposure of female mice to TBT, inorganic arsenic and total western diet lead to sexually dimorphic metabolic disruption in their offspring

At 8 weeks of age and after 3.5 weeks of exposure, we mated F0 exposed females with same-age unexposed males for a week to produce their F1 progeny (Figures 1A and 2A). At F1 weaning, we selected at least 10 mice for each sex and exposure group to determine if ancestral exposures resulted in F1 metabolic disruptions (Figure 2A). We tracked the body weight of F1 mice weekly since weaning at 3 weeks of age until we euthanized them at 11 weeks of age (Figure 2A). In the past, we demonstrated that exposure to secondary metabolic challenges, such as a high-fat diet, could reveal metabolic disruptions caused by ancestral exposures^8,10^. We fed F1 mice CD since weaning until 7 weeks of age and TWD from that point until the end of the experiment (Figure 2A). At 7 weeks of age, when we changed diets, we fasted F1 mice for 12 hours and measured their body weight and glucose levels in peripheral blood (Figure 2A). At 11 weeks of age, we fasted F1 mice for 12 hours before euthanizing them and measuring their body weight, glucose levels in peripheral blood, and the weight of gonadal and inguinal white adipose tissue depots (gWAT, and iWAT, respectively) and liver (Figure 2A). We euthanized the animals by isoflurane overdose and cardiac exsanguination to harvest cardiac blood, gWAT, iWAT and liver (Figure 2A). We processed cardiac blood samples obtained during euthanasia to determine levels of 11 relevant metabolic regulators, namely amylin, GIP, ghrelin, GLP-1, insulin, leptin, PYY, glucagon, PP, resistin and C-peptide (Eve Technologies Mouse, Rat Metabolic Array (MRDEMT)).

**Figure 2.**
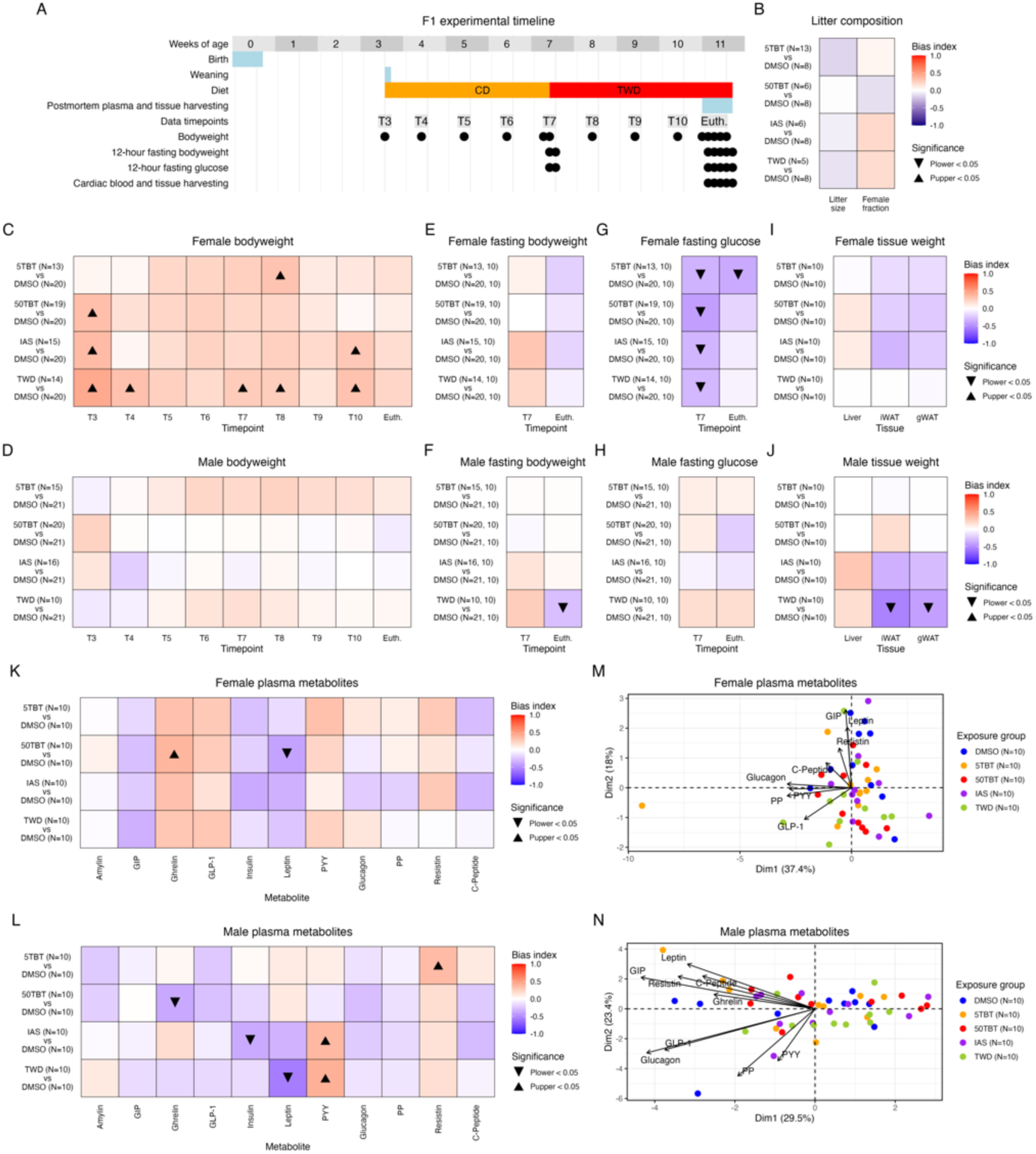
F1 offspring metabolic effect of the preconception exposure of female mice to TBT, inorganic arsenic, and Total Western Diet. **A.** Experimental timeline detailing primary mouse operations and the timepoint structure employed to assess variations in litter size and sex ratio, body weight, fasting body weight, fasting glucose, tissue weight, and plasma metabolite levels resulting from maternal preconception exposure to TBT, IAS, and TWD when compared to controls. **B.** Comparison of litter size (N) and sex ratio (female fraction) between exposure groups and controls using uMCW tests. **C-D.** Comparison of body weight normalized by age (g/days old) between exposure groups and controls using uMCW tests for F1 females and males, respectively. **E-F.** Comparison of fasting body weight normalized by age (g/days old) between exposure groups and controls using mbMCW tests for F1 females and males, respectively. **G-H.** Comparison of fasting glucose normalized by body weight and age ((mg/dL)/g/days old) between exposure groups and controls using uMCW tests for F1 females and males, respectively. **I-J.** Comparison of tissue weight normalized by age (g/days old) between exposure groups and controls using uMCW tests for F1 females and males, respectively. **K-L.** Comparison of metabolite plasma level normalized by body weight and age ((pg/mL)/g/days old) between exposure groups and controls using uMCW tests for F1 females and males, respectively. **M-N.** Principal component analyses (PCAs) of metabolite plasma levels normalized by body weight and age (pg/mL/g/days old) for females and males in exposure and control groups, respectively. The number of replicates (N) in each experimental group is indicated in each panel.

We used uMCW tests to determine whether mouse body weights, fasting glucose, tissue weights, plasma metabolite levels, and litter size and sex ratio were significantly biased in the same direction for each exposure group when compared with controls (Figure 2B-L). To determine if the F1 groups were metabolically distinctive, we performed principal component analyses (PCA) using the plasma metabolite data (Figure 2M-N). To maximize the value of biological replication, we conducted PCA analyses using only those metabolites whose plasma levels had been successfully determined in all biological replicates for all groups in each sex (8 in females and 9 in males).

We did not find any significant differences in litter size or sex ratio between the exposure groups and the controls (Figure 2B). This suggests that the exposures did not cause any obvious negative effects on the fertility of the F0 females or the viability of their F1 offspring.

Upon analyzing all the metabolic traits, distinct differences were observed between the sexes. Female groups of all treatments exhibited comparable signs of metabolic disruption compared to controls. They exhibited increased body weight, particularly before weaning and after the diet change at 7 weeks of age (Figure 2C). Additionally, they demonstrated lower plasma glucose levels, particularly before the diet change (Figure 2G). Furthermore, similar alterations were observed in GIP, ghrelin, GLP-1, insulin, leptin, and PYY (Figure 2K). While other changes were not statistically significant or as pronounced as the preceding ones, the F1 females from the exposure groups exhibited similar trends compared to controls. For instance, they exhibited higher and lower fasting body weight before the diet change and euthanasia, respectively (Figure 2E). They also experienced a reduction in gWAT and iWAT compared to controls (Figure 2I). Although not all the trends or significant alterations were precisely identical for the exposure groups, particularly the 5TBT group, the similarities between them were more pronounced than the differences.

In contrast, F1 male groups exhibited less pronounced and more disparate signs of metabolic disruption compared to controls. TWD mice exhibited more evident metabolic disruption, characterized by lower fasting body weight, gWAT, and iWAT weights, and lower and higher plasma levels of leptin and PYY, respectively, at euthanasia (Figure 2F, J and L). Notably, TWD male mice were the sole group exhibiting moderate differentiation from control male mice when analyzing plasma levels of nine metabolites using PCA (Figure 2N). IAS F1 males exhibited significantly higher and lower plasma levels of PYY and insulin, respectively (Figure 2L). IAS also exhibited non-significant lower trends in gWAT and iWAT weights and plasma levels of leptin at euthanasia, resembling significant alterations observed in the TWD group (Figure 2J, 2L). The two TBT groups exhibited less pronounced metabolic disruption. The only significant alterations in the 5TBT and 50TBT groups were in the plasma levels of resistin and ghrelin, respectively (Figure 2L).

### Preconception exposure of female mice to TBT, inorganic arsenic and total western diet lead to alterations of the expression of mitochondrial genes in their offspring

We processed gWAT and liver samples to perform transcriptomic analyses because they are relevant tissues for regulation of energy homeostasis, and because we have previously demonstrated that transcriptomic analysis of gWAT and liver can be instrumental in substantiating the mechanisms underlying the multigenerational metabolic disruptions induced by environmental exposures^8,9^. A general observation of the expression patterns of the 100,000 most expressed genes across all samples show that their transcriptome is clearly distinctive between tissues and sexes, the latter distinction being more pronounced in gWAT than in liver (Figure 3A).

**Figure 3.**
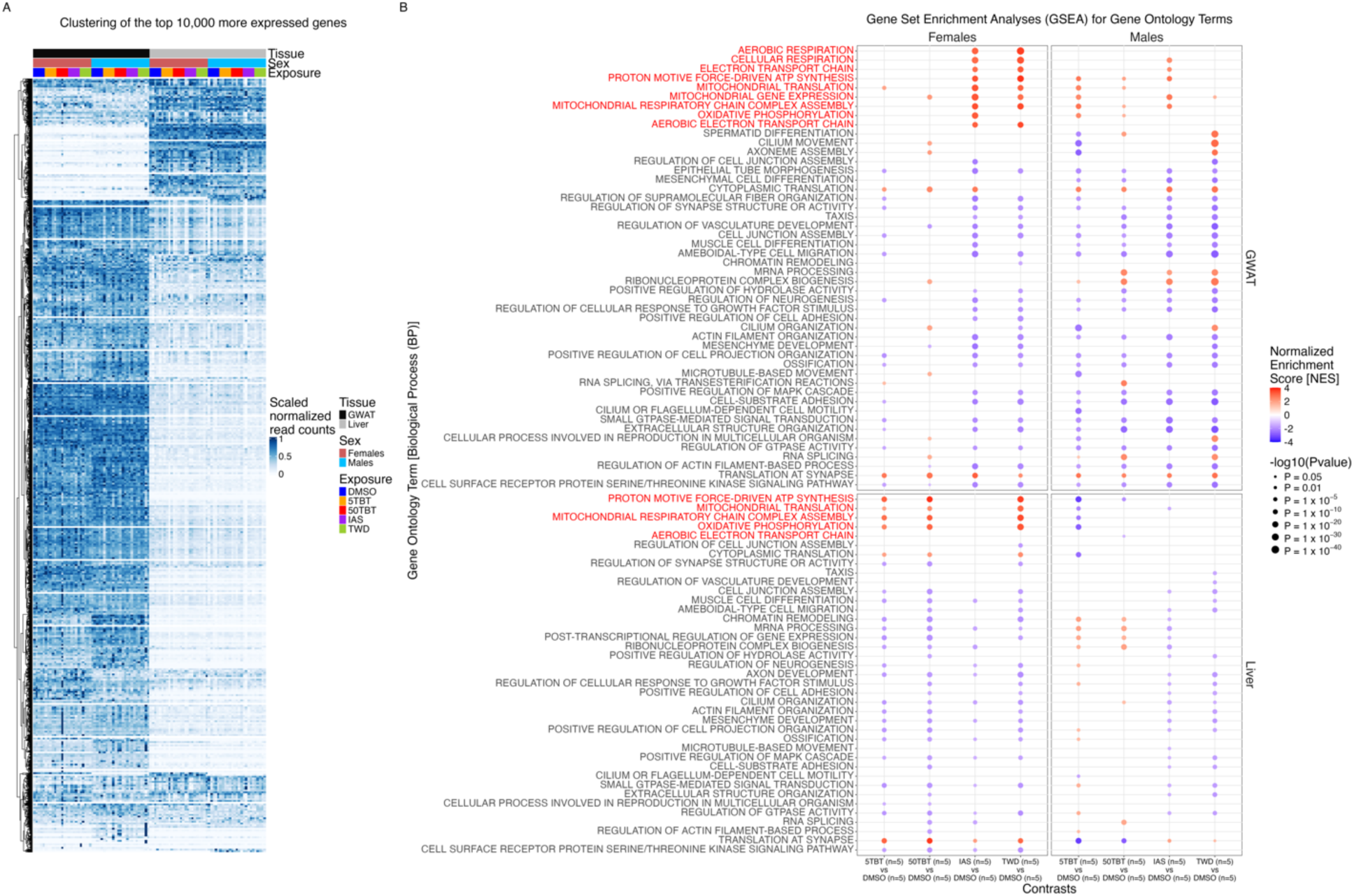
F1 offspring transcriptomic effect of the preconception exposure of female mice to TBT, inorganic arsenic, and Total Western Diet. **A.** Gene expression patterns across 100 RNA samples (2 sexes x 2 tissues x 5 experimental groups x 5 replicates) for the 100,000 more expressed genes. **B.** Results of Gene Set Enrichment Analysis (GSEA) using Gene Ontology (GO) terms of the Biological Process (BP) categories for all exposure group *versus* control contrasts. For each gene and contrast, we conducted uMCW tests to assess the significance of the expression difference between exposure groups and controls. Genes were ranked for each contrast using uMCW test BIs from highest to lowest, and GSEA analysis was performed using the R package *fgsea*. In this analysis, we present GSEA results for the 50 GO-BP with more extreme GSEA *P* values across contrasts. GO-BP terms marked in red labels at the top of each panel correspond to metabolic GO-BP terms. The number of replicates (N) in each experimental group is indicated in the *x*-axis.

To elucidate gene expression alterations and their potential functional implications, we conducted uMCW tests for each gene across each of the 16 exposure group *versus* control contrasts (4 exposure groups x 2 tissues x 2 sexes). Subsequently, we performed Gene Set Enrichment Analyses (GSEA) using Gene Ontology (GO) terms from the Biological Process (BP) category. Figure 3B presents GSEA results for GO-BP terms with the 50 highest -log10(P values) across all 16 exposure *versus* control contrasts. These findings demonstrate evident metabolic disruptions, consistent with a perturbation of mitochondrial functions, with distinct differences observed between exposure groups and sexes. In females, mitochondrial GO-BP terms exhibit significant overrepresentation for genes with higher expression in gWAT in IAS and TWD groups compared to controls. Additionally, these terms are overrepresented for genes exhibiting higher expression in liver in 5TBT, 50TBT, and TWD groups relative to controls. Conversely, in male subjects, mitochondrial GO-BP terms are overrepresented for genes with higher expression in gWAT in 5TBT and IAS groups compared to controls, and for genes with lower expression in 5TBT compared to controls.

The observed enrichment patterns in GO-BP suggest that the similar metabolic disruption observed in F1 female exposure groups (Figure 2) may be dependent on alterations in mitochondrial function (Figure 3B). However, there are also relevant distinctions. For TBT groups, the mitochondrial alterations appear to affect the liver more significantly than gWAT. In contrast, for the IAS group, gWAT appears to be the tissue most affected by the mitochondrial alterations. Finally, for the TWD group, mitochondrial functions seem to be altered in both tissues.

The relationship between metabolic and transcriptomic alterations in males is less clear. The TWD group appears to exhibit more distinctive metabolic alterations when compared to controls (Figure 2), and it also presents clear functional alterations, primarily in gWAT, but not related to mitochondrial function (Figure 3B). Therefore, the potential relationship between these two alterations does not appear straightforward. Conversely, the 5TBT and IAS groups, which exhibited limited signs of metabolic disruption (Figure 2), do demonstrate potential alterations in mitochondrial functions (Figure 3B).

### Preconception exposure of female mice to TBT, inorganic arsenic and total western diet lead to sexually dimorphic alterations of HEC in their offspring

We and others have demonstrated that transcriptomic analyses, guided by indirect approximations of chromatin organization, serve as valuable tools to identify preliminary evidence suggesting the potential of environmental exposures to induce chromatin organization perturbations in the descendants of exposed individuals^8,9,13^. A hallmark of all these cases was a dichotomous pattern of significant changes in gene expression for genes located in regions that approximated the heterochromatic and euchromatic compartments. The expression of genes located in heterochromatin was significantly biased in one direction, while the expression of genes located in euchromatin was significantly biased in the other direction^5,6,10^.

Our indirect approach to ascertain HEC alterations through transcriptomic, DNA methylation, and chromatin accessibility data employed the utilization of isochores^8,9^. Isochores are long segments of chromosomes characterized by a pronounced propensity for base composition uniformity, typically categorized into five classes: L1, L2, H1, H2, and H3, ranging from the most AT-rich to the most GC-rich^61,62^. The relevance of isochores as HEC approximations for integrative analyses stems from the following considerations: (i) base composition is invariant across sexes, developmental stages, cell types, or individuals within the same species; (ii) individual isochores approximate well local structural motifs of chromatin organization, such as topologically associating domains (TADs)^61,62^; and (iii) AT- and GC-rich isochores effectively represent the heterochromatin and euchromatin compartments of eukaryotic genomes^52,53^. Specifically, AT-rich L1 and L2 isochores correspond closely to the heterochromatic compartment, while GC-rich H2 and H3 isochores effectively represent the euchromatic compartment^61,62^.

To ascertain whether female mice preconception exposure to TBT, IAS, and TWD induced coordinated alterations of gene expression for genes overlapping isochores, we conducted biased-measures MCW (bMCW) tests. The bMCW test is a combination of two tests that assess whether a set of measures of bias for a quantitative trait between two conditions and a specific subset of these bias measures are themselves significantly biased in the same direction. We used bMCW tests to determine whether gene expression differences between each exposure group and controls for genes in the whole transcriptome, overlapping specific isochores, and overlapping isochores of the same class or located within autosomes and the *X* chromosome were significantly biased in the same direction. The measure for gene expression difference we used for bMCW tests was the uMCW test BIs obtained for each gene and exposure group *versus* control contrast. The goal of the analysis of concerted biases of expression for the whole transcriptome and at the chromosome level was to provide context to the results obtained using isochores. Given that in vertebrates all chromosomes possess isochores belonging to the five classes^36,63,64^, chromosomes represent an independent yet interrelated motive of chromatin organization distinct from the motives approximated by isochore classes.

Our isochore-based bMCW test results exhibited distinct differences between sexes that are comparable to those observed in other metabolic traits (Figure 2). Specifically, all female groups exhibited similar patterns, while in the male groups, TWD stood out as the most distinctive from the other three.

In female gWAT, the transcriptome exhibits a general tendency to be overexpressed in 50TBT males and underexpressed in the other three groups compared to controls (Figure 4B). Despite these disparities, the significance of the expression biases for genes in AT- and GC-rich isochores reveals a similar dichotomous pattern across all four groups (Figure 4C). The bias in expression for genes situated in GC-rich isochores is higher than what would be expected by chance, while the bias in expression for genes in AT-rich isochores is lower than what would be expected by chance. In female liver, the patterns between groups are even more uniform. The transcriptome as a whole is underexpressed when compared to controls (Figure 4B). Additionally, the significance of the expression biases for genes in AT- and GC-rich isochores demonstrates the same dichotomous pattern between groups, closely resembling the pattern observed in gWAT (Figure 4C).

**Figure 4.**
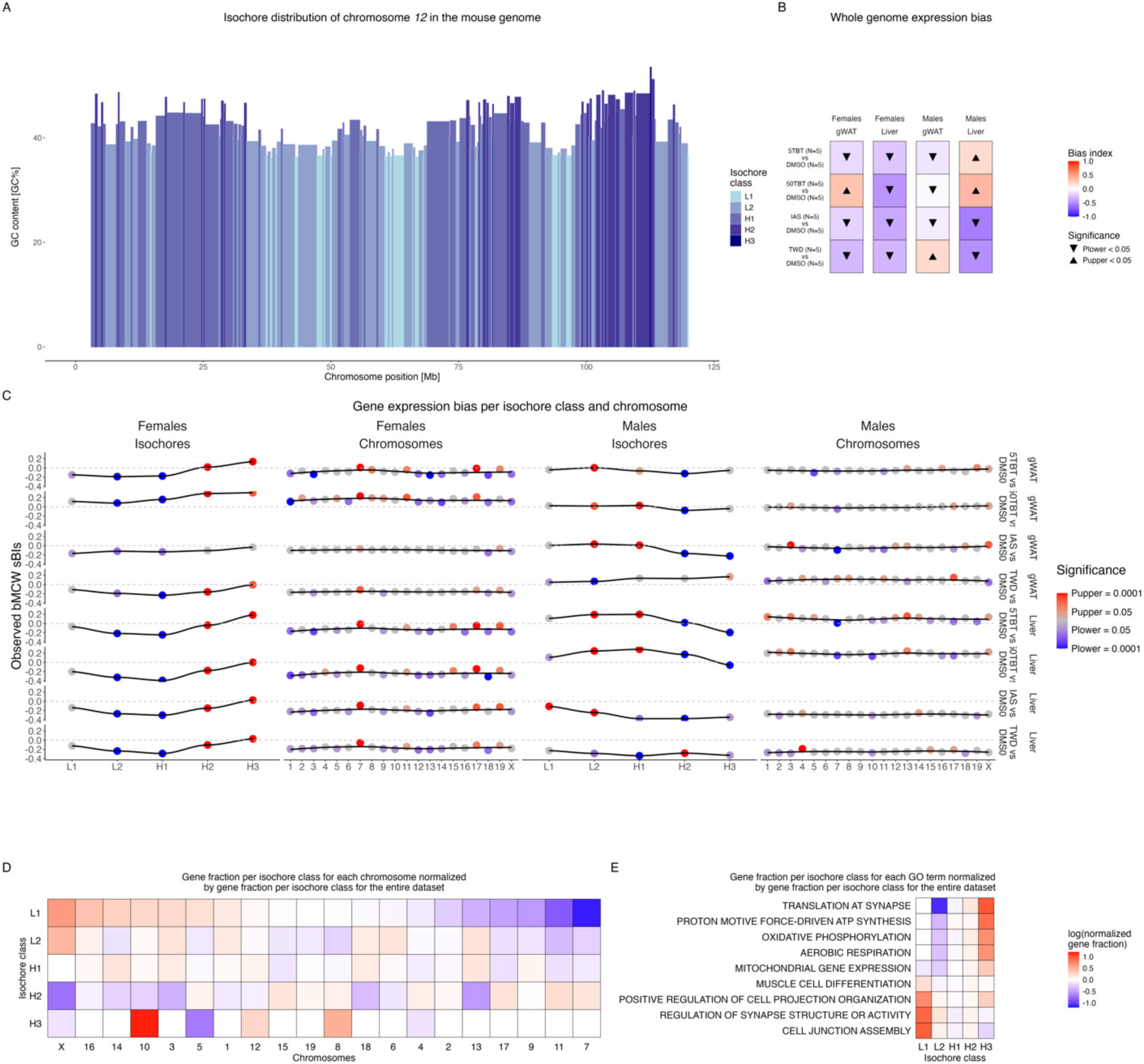
Isochore distribution of F1 transcriptomic alterations caused by the preconception exposure of female mice to TBT, inorganic arsenic, and Total Western Diet. **A.** Isochore composition of chromosome *12* in the mouse genome. Isochore classes, L1, L2, H1, H2, and H3, are defined by their GC content, ranging from the most AT-rich to the most GC-rich. **B.** Analyses of concerted alterations in gene expression for all genes in the transcriptome and each exposure group *versus* control contrast were using bMCW tests. **C.** Analyses of concerted alterations in gene expression for genes located within each of the five isochore classes and each autosome and the *X* chromosome for female and male, gWAT and liver samples for all exposure groups *versus* control contrasts using bMCW tests. **D.** Relative gene distribution within each isochore class for all autosomes and the *X* chromosome for genes in the female gWAT 50TBT *versus* control contrast. **E.** Relative gene distribution within each isochore class for 10 GO-BP terms selected from the results in Figure 3B for genes in the female gWAT 50TBT *versus* control contrast. bMCW sBIs: biased Monte Carlo-Wilcoxon test subset-Bias Indexes (see Methods).

In male gWAT, groups 5TBT, 50TBT, and IAS exhibit similar patterns. The transcriptome as a whole tends to be underexpressed when compared to controls (Figure 4B). Furthermore, the significance of the biases in expression for genes in AT- and GC-rich isochores demonstrates the same dichotomous patterns, namely that gene expression bias for AT- and GC-rich isochores are higher and lower than what would be expected by chance, respectively (Figure 4C). These last patterns are the exact opposite of the ones observed for the females of the same groups in gWAT (Figure 4C). The patterns observed in male TWD gWAT clearly differ from the other three groups (Figure 4C).

In male liver, the heterogeneity between groups is more pronounced. In the TBT groups, the transcriptome as a whole tends to be overexpressed when compared to controls (Figure 4C). Additionally, the significance of the biases in expression for genes in AT- and GC-rich isochores is precisely the same as in male gWAT and the exact opposite of female liver of TBT groups (Figure 4C). In the case of IAS and TWD groups, the transcriptome as a whole tends to be underexpressed when compared to controls (Figure 4B). Regarding the significance of the biases in expression for genes in AT- and GC-rich isochores, the IAS group resembles more the patterns observed for TBT groups than the TWD group does (Figure 4C).

The chromosome-based bMCW tests also yield significant results, albeit with less pronounced bias values compared to their isochore-based bMCW test counterparts (Figure 4C). In fact, the results obtained for chromosomes can be attributed to their distinct isochore composition and the biases observed in each isochore class (Figure 4C-D). For instance, the expression of genes in chromosomes *7* and *11*, which contain more GC- than AT-rich isochores (Figure 4D), tend to be biased in the same direction as GC-rich isochores do for each exposure *versus* control contrast (Figure 4C). Conversely, *X* and *3* chromosomes, which contain more AT- than GC-rich isochores (Figure 4D), tend to be biased in the same direction as AT-rich isochores do for each exposure *versus* control contrast (Figure 4C).

The dichotomous patterns of gene expression bias observed in AT- and GC-rich isochores align with our hypothesis that preconception exposure to TBT and IAS may have resulted in sexually dimorphic HEC perturbations in their offspring. Conversely, while our findings for TWD also support HEC perturbations in the offspring of exposed females, such perturbations appear to be non-sexually dimorphic.

To determine whether such perturbations could be interrelated with the functional alterations previously observed (Figure 3B), we analyzed the isochore distribution of genes associated with some of the GO-BP with more pronounced results (Figure 3B). Figure 4E illustrates the isochore distribution of genes associated with five GO-BP terms overrepresented in genes that were underexpressed in the exposure groups compared to controls, and five GO-BP terms overrepresented in genes that were overexpressed in the exposure groups (Figure 3B). Notably, the two sets of terms exhibit contrasting bias regarding the gene isochore distributions. Genes associated to GO-BP terms that were overrepresented for genes that were overexpressed in the exposure groups tend to be preferentially located in GC-rich isochores, while genes associated to GO-BP terms that were overrepresented for genes that were underexpressed in the exposure groups tend to be preferentially located in AT-rich isochores.

These latter results suggest a potential interplay between HEC and functional perturbations induced by maternal preconception exposure to TBT, IAS, and TWD in their offspring’s gWAT and liver. To further explore this interplay, we compared gene expression biases for genes belonging to two large gene families with broad but distinct isochore distributions, the specific isochores where each member of these families is located, and the five isochore classes in gWAT and liver for all the exposure *versus* control contrasts (Figure 5).

**Figure 5.**
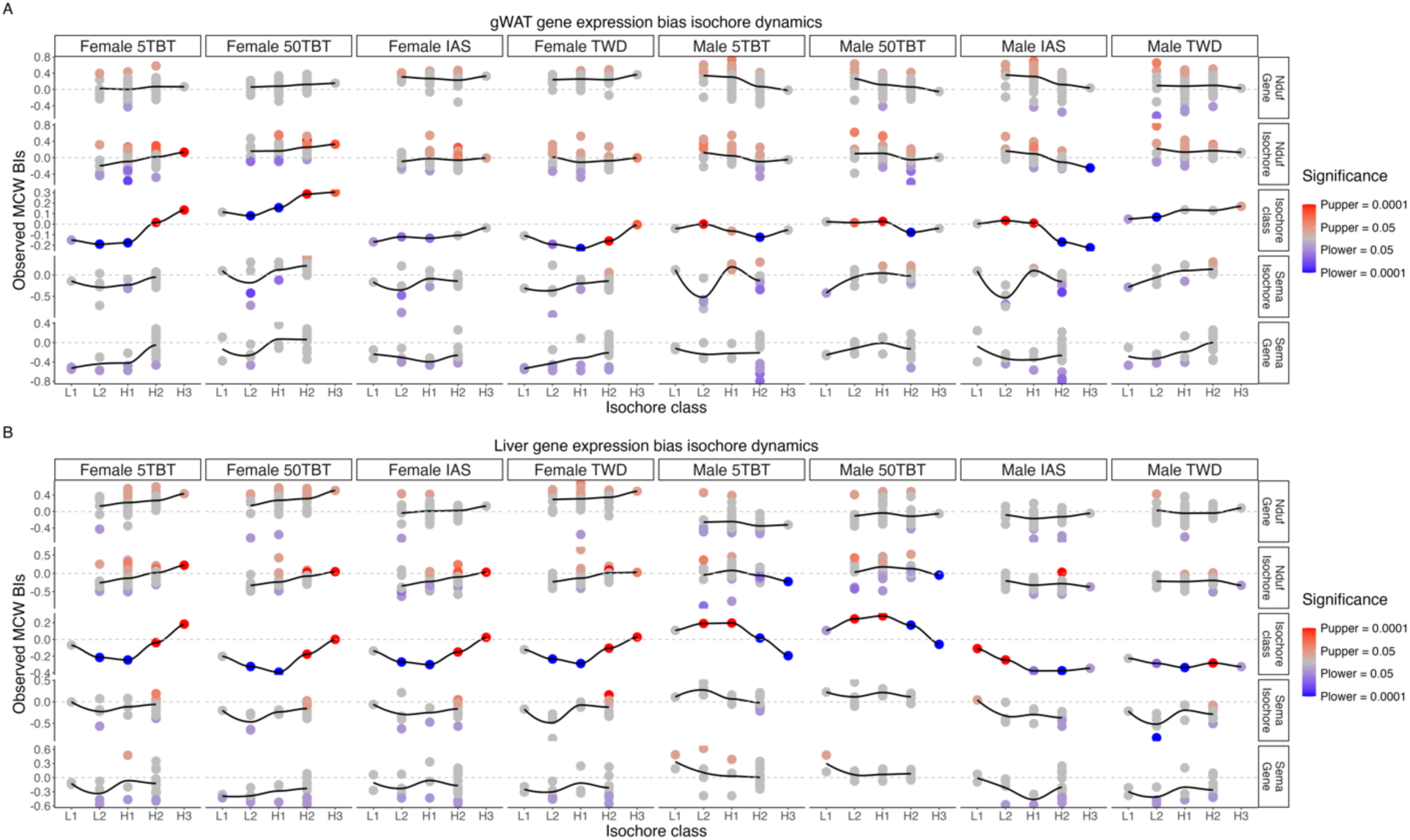
Concerted changes in gene expression caused by the preconception exposure of female mice to TBT, inorganic arsenic, and a total Western diet at three levels of resolution in the mouse genome. Analysis of gene expression bias for genes belonging to the Nduf and Sema gene families, the isochores where each of these genes reside, and each of the isochore classes for each of the exposure group *versus* control contrasts in gWAT (**A**) and liver (**B**). Gene expression bias analyses for individual genes were conducted using uMCW tests. Gene expression bias analyses for specific isochores and isochore classes were performed using bMCW tests. Gene expression bias anlyses results for isochore classes results are also represented in Figure 4C. MCW BIs: Monte Carlo-Wilcoxon test Bias Indexes (see Methods).

We initially identified the Sema and Nduf gene families due to their distinct association with GO- BP terms overrepresented in over- and underexpressed genes across all the exposure *versus* control contrasts. The mitochondrial Complex I (NADH:ubiquinone oxidoreductase) is a protein complex that participates in the respiratory chain, with subunits encoded both in the mitochondrial and nuclear genomes^65^. The Nduf gene family comprises all genes encoding nuclear subunits of Complex I. Nduf genes were associated with all the mitochondrial GO-BP terms predominantly overrepresented in genes that were overexpressed in exposure groups when compared with controls (Figure 3B). The Sema gene family encodes Semaphorins, a large family of surface receptors and ligands that regulate multiple processes in neuronal, vascular, immune, bone, and epithelial systems^66,67^. Sema genes were associated with diverse GO-BP terms such as “*Ameboidal type cell migration*,” “*Axon development*”, “*Mesenchyme development*,” “*Regulation of neurogenesis*” and “*Taxis*,” which were mostly overrepresented in genes that were underexpressed in exposure groups when compared with controls (Figure 3B).

Our transcriptomic dataset comprises 55 Nduf genes (including three pseudogenes) and 20 Sema genes. While both gene families are represented in four out of the five isochore classes, the Nduf isochore distribution exhibits a slight preference for GC-rich isochores, whereas the Sema isochore distribution exhibits a slight preference for AT-rich isochores (Figure 5).

To ascertain whether the expression of genes and groups of genes located within the same isochore mirrored the patterns observed for isochore classes, we identified the isochores in which each Nduf and Sema gene were located. Subsequently, we conducted bMCW tests to determine if the expression of the genes contained within those isochores was altered in a concordant manner. Figure 5 presents the outcomes of bMCW tests for each isochore class (also represented in Figure 4), bMCW tests for specific isochores harboring Nduf or Sema genes, and uMCW tests for each Nduf and Sema gene.

Notably, despite certain exceptions, the expression of individual Nduf and Sema genes and the specific isochores in which they reside appear to correspond with the dichotomous patterns observed for the five isochore classes. For instance, in female 5TBT liver, genes situated within GC-rich H2 and H3 isochores tend to be overexpressed, while genes located within AT-rich L1, L2 and H1 isochores tend to be underexpressed relative to controls (Figure 5B). This dichotomy is replicated with varying degrees of intensity for Nduf and Sema genes and the specific isochores where each of these genes is situated.

In summary, our isochore-based analyses support the hypothesis that preconception exposure to TBT, IAS, and TWD results in HEC perturbations in their offspring, which are perceived at varying levels of resolution: specific genes, localized regions encompassing multiple genes defined by their base composition uniformity, and dispersed genomic regions characterized by similar base composition.

### Preconception exposure of female mice to TBT, inorganic arsenic and total western diet might alter the chromatin architecture regulating the expression of leptin in females but not in males

To delve deeper into the association between the putative disruptions we observed in the descendants of female mice exposed to TBT, IAS, and TWD prior to conception for metabolic traits and chromatin organization, we redirected our focus to the study of the gene *Lep*, which codes for the hormone leptin. Leptin, primarily secreted by adipose tissue, plays a crucial role in regulating feeding behaviors by informing the hypothalamus of the state of the energy reserves stored in adipose tissue^68,69^. When these reserves are full, leptin secretion increases, reason why leptin is often referred to as the “satiety hormone”. For example, leptin levels in plasma increase during meals, to inform the brain about satiety and prevent the organism from overfeeding. Recent research has proposed that a chromosome loop structure involving neighboring genes *Lnclep* and *Lep* is pivotal for regulating the expression of the leptin gene, and that the expression of *Lnclep* and *Lep* exhibits a strong correlation across various pathophysiological conditions^70–72^.

We previously noted that plasma levels of leptin exhibited varying degrees of decrease in F1 females belonging to the four exposure groups and in F1 males for the IAS and TWD groups (Figure 2K-L). Furthermore, leptin plasma levels appeared to be a significant metabolite contributing to the PCA separation of TBT, IAS, and TWD groups from controls in females and TWD from the other groups in males (Figure 2M-N). To elucidate whether female preconception exposure to TBT, IAS, and TWD induces alterations in the chromatin organization of the gene *Lep*, which may explain the observed changes in leptin plasma levels, we started by identifying the isochore where the gene *Lep* is situated.

Figure 6A shows the results of our uMCW tests for all genes located within the same H2 isochore, encompassing the *Lep* and *Lnclep* genes, for the 16 exposure *versus* control contrasts. Notably, both the *Lep* and *Lnclep* genes are absent from liver tissue, consistent with the well-established adipose-specificity of the *Lep* gene^73^.

**Figure 6.**
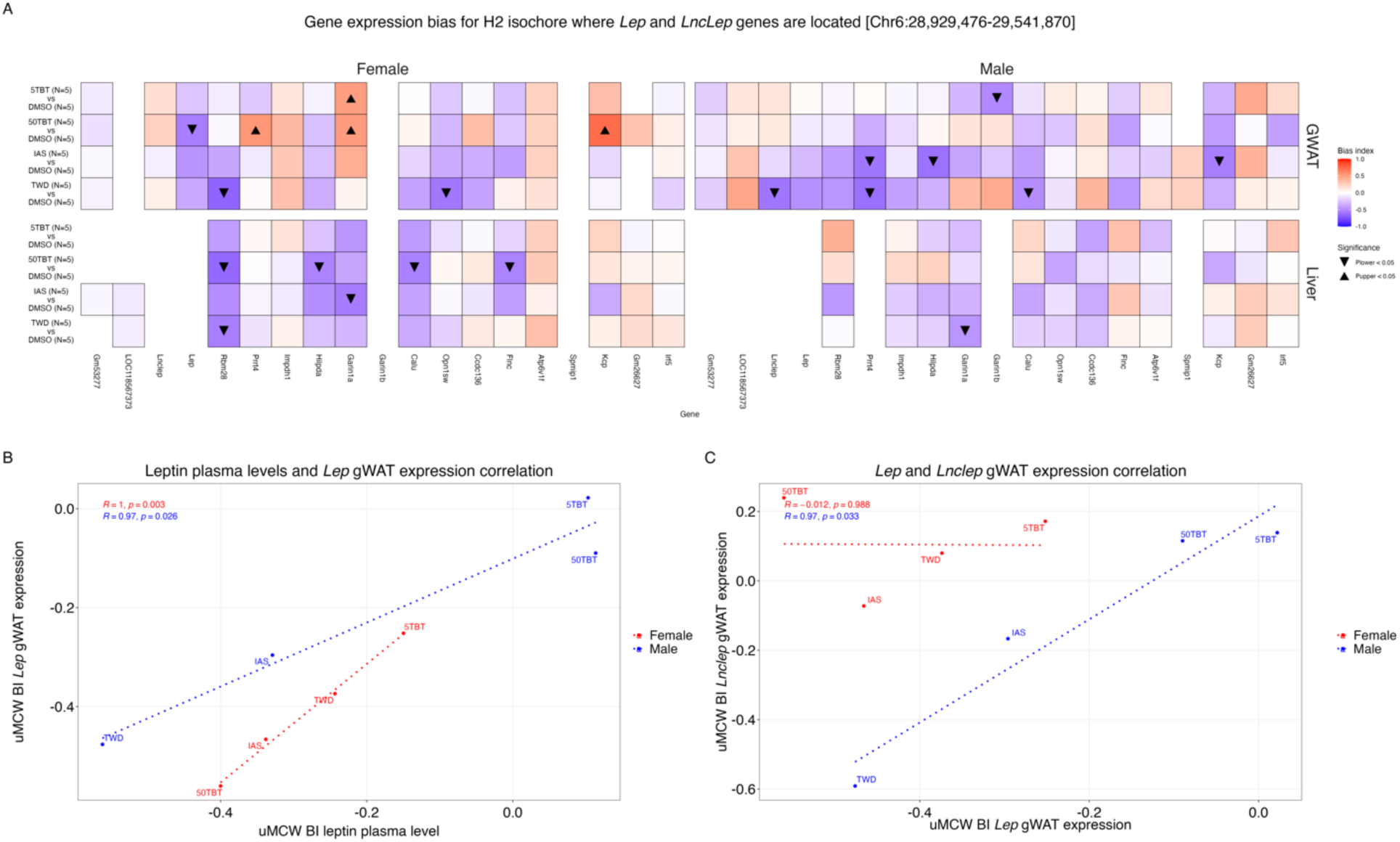
Concerted changes for leptin caused by the preconception exposure of female mice to TBT, inorganic arsenic, and a total Western diet. **A.** Analyses of gene expression bias for genes in the H2 isochore where the gene *Lep* is located conducted using uMCW tests. Missing tiles represent cases for which a gene was deemed non-expressed (see Methods). **B.** Pearson correlation analyses between changes in leptin plasma levels and Lep gene expression in gWAT for female and male exposure groups *versus* control contrasts determined using uMCW tests. **C.** Pearson correlation analyses between changes in the expression of *Lep* and *LncLep* genes in gWAT for female and male exposure groups *versus* control contrasts determined using uMCW tests. uMCW BI: unmatched-measures Monte Carlo-Wilcoxon Bias Index.

Figure 6B shows the highly statistically significant correlations between the decreases in plasma levels of leptin and the expression of the *Lep* gene for both female and male exposure *versus* control group. Given that the primary tissue responsible for leptin secretion is adipose tissue, these findings suggest that the declines in plasma leptin observed in the descendants of females exposed to TBT, IAS, and TWD are attributed to alterations that diminish the expression of the *Lep* gene within the adipose tissue. Notably, our results demonstrating the absence of *Lep* expression in liver tissue and the precise correlations between variations in plasma leptin and the expression of the *Lep* gene serve as validation of our analytical approach in effectively capturing the adipose-specificity of the *Lep* gene.

Figure 6C shows the correlation between the results of uMCW tests for the expression of *Lep* and *Lnclep* genes for female and male exposure *versus* control contrasts separately. In this case, there is an almost perfect correlation between the changes in expression for both genes in males, which would agree with afore mentioned literature^70–72^. However, in females the changes in expression for both genes are not at all correlated. These results would suggest that although in males the regulatory context of *Lep* and *Lnclep* that mediates the correlation of their expression in physiological conditions is preserved, in females such context is perturbed.

## Discussion

In this study, we investigated whether preconception exposure of female mice to three metabolic disruptors induced metabolic alterations in their offspring that could be mediated by perturbations of chromatin organization. Our findings generally align with this scenario, while simultaneously presenting some unexpected results. If we were solely presented with the effects observed in exposed females and their male offspring, we would conclude that each of the exposure studied here exhibited different effects, with TWD exposure being the most pronounced and distinctive.

Contrary to the direct and male offspring effects, the results observed for female offspring are surprising in two directions. Firstly, the phenotypes observed for the four experimental groups when compared to controls exhibited more similarities than differences (Figures 2-5). Secondly, the perturbations we detected in offspring females suggest they exhibit a phenotype consistent with atypical anorexia nervosa in human populations. We observed low levels of plasma leptin, insulin, and glucose, high levels of plasma ghrelin, GLP-1, and PYY, and low fractions of adipose tissue in TBT, IAS, and TWD females when compared with controls (Figure 2). Although many of these alterations would not be considered significant with the commonly used threshold of significance (*P* = 0.05), it is remarkable that all such trends have been observed in anorexic human females^74–79^. Furthermore, altered mitochondrial functions, such as those we observed in gWAT and liver transcriptomic analyses (Figure 3), have been reported in studies of transcriptome variation between anorexic patients and healthy controls^80^.

Although these trends align with anorexic phenotypes, it is challenging to definitively determine whether female offspring of female mice exposed to TBT, IAs, or TWD exhibit anorexia, especially considering that self-starvation and substantial weight loss are the primary diagnostic characteristics associated with such condition^81^. Firstly, we lack valid data on the feeding behavior of offspring females in our study. As per the experimental design, we did not conduct individualized measurements of food consumption, and the number of cage replicates per experimental group was limited. Secondly, in response to a fasting challenge, we did not observe significant nor consistent variation trends for TBT, IAs, or TWD females when compared to controls (Figure 2E). Thirdly, females belonging to the TBT, IAS, and TWD groups exhibited a tendency to be overweighted when compared to controls, with varying degrees of significance across the study period (Figure 2C). Atypical anorexia nervosa is used to describe patients who exhibit signs of anorexia but do not experience a decrease in body weight^81^. While further characterization of the offspring effects induced by preconception exposure to metabolism disruptors is warranted, it is intriguing to speculate that such exposures may predispose to atypical anorexia nervosa. Despite the growing recognition of the significance of metabolic alterations in the pathogenesis and manifestation of anorexia nervosa^82,83^, and the availability of numerous animal models for the study of anorexia nervosa^84^, to the best of our knowledge, no empirical evidence has been found establishing a causal relationship between anorexic phenotypes and exposure to metabolism disruptors.

A distinctive characteristic of our study is its high degree of integration utilizing disparate data. Our study yielded data spanning multiple metabolic traits, both basally and in response to metabolic challenges such as fasting or diverse diets, as well as transcriptomic data from two generations, two F1 sexes, and two tissues with distinct ontogenies. The integration was facilitated by three key factors.

Firstly, a standardized methodology was employed to discern differences between exposure groups and controls for all traits. The fact that various MCW tests, each interrogating different data structures and traits, yielded comparable metrics such as BIs, *P_upper_*, and *P_lower_* significantly contributed to establishing connections between different traits. A validation of this point is the substantial correlation observed between plasma levels of leptin and gene expression in gWAT. Another point of validation is that results obtained for individual metabolites closely resemble those presented in PCAs.

Secondly, we used non-stringent definitions of statistical significance. While our study aimed to find evidence consistent with a hypothesis, it is inherently exploratory in nature. Although special attention is often given to distinguishing true and false positives, such as through multiple test corrections, it is equally perilous for exploratory studies to discard potentially true positives by employing a highly stringent threshold of significance, which could result in false negatives. To determine potentially significant results, we leveraged the integration of multidimensional data. As we just mentioned, such integration was facilitated by employing the same analytical approach, MCW tests, but also by employing distinct approaches to measure the same trait at different levels, *e.g.*, leptin plasma levels and gWAT expression (Figure 5) or gene expression biases for specific genes, the isochores where they are located, and isochores of the same class dispersed throughout the genome (Figure 4). In the case of plasma levels of metabolites, we observed similar patterns in females even when these metabolites were not considered significant using conventional non-stringent definition of significance (*P* = 0.05), that resemble the weight of such metabolites in PCAs (Figure 2).

Lastly, although we did not intend originally to study exposures that were expected to yield highly similar outcomes in the female offspring of exposed females, this fact contributed to the identification of potentially relevant results. As previously indicated, the similar behaviors of plasma metabolites or body weight in the four exposure groups when compared with controls appeared highly relevant to entertain the possibility that female preconception exposure to metabolic disruptors can predispose to sexually dimorphic atypical anorexia nervosa regardless of the number of data points that could be considered significant (*P* < 0.05).

Despite these strengths, this study also has limitations. The primary objective of our study was to employ methodologies that are widely accessible to laboratory researchers. These included standard balances or measure cylinders for measuring body weight, food and water consumption, facility-serviced bulk transcriptomic analyses to assess gene expression, and publicly available computer tools for integrative analyses. The intentionally low resolution of our study caused that our limited ability to capture feeding behaviors that reinforce the anorexic state was not the sole limitation.

For example, the central *a priori* hypothesis of our study posited that female preconception exposure to metabolic disruptors could lead to offspring metabolic alterations mediated by changes in the germ line of exposed individuals. These changes, in turn, could affect the offspring’s establishment of chromatin organization, which has the potential to self-propagate throughout development and predispose to metabolic disruption. However, we cannot entirely rule out the possibility that the effects we observed were influenced by the direct action of exposure agents, such as TBT, IAS, or components of TWD, or the results of their metabolization, like dibutyltin (DBT) in the case of TBT^85^, lingered or bioaccumulated in the mother after the interruption of the exposure immediately prior to fertilization on the developing embryo. Additionally, the influence of alterations caused by the exposures in the mother during the development of the embryo could also contribute to the effects we aimed to study and their direction. Future studies based on the *in vitro* fertilization of oocytes from exposed females are necessary to disentangle the potential contribution of the effects of environmental exposure on germ cells of the exposed individual, the developing embryo, and/or the physiological alterations of the mother during gestation.

Furthermore, our analyses to ascertain offspring disruptions in metabolism and chromatin organization are neither exhaustive nor direct. In the case of metabolic disruptions, a larger and more intricate battery of metabolic determinations would be necessary to comprehensively comprehend the nature of the metabolic disruptions that our study suggests may have occurred. Similarly, the results of our isochore-based approach can only be regarded as indirect evidence of such perturbations. Although methodologies exist to directly interrogate chromatin organization, such as high-throughput chromosome conformation capture (Hi-C), their application in the context of complex studies and tissues remains quite limited^86^. Consequently, we view this study as the preliminary work required to establish whether there is potential for conducting more comprehensive studies on the offspring effects of exposure to metabolic disruptors.

## Conclusion

This study provides support for the hypothesis that preconception exposure to metabolic disruptors can cause metabolic alterations in the offspring of exposed individuals through perturbations of chromatin organization. Although our study has the low analytical resolution typical of preliminary exploratory studies, it is noteworthy the level of congruence observed after integrating multidimensional data. This validates our approach as a preliminary step towards determining the potential of environmental exposures to lead to multigenerational effects. Beyond the methodological value of our study, the potential connection we uncovered between preconception exposure to metabolic disruptors and atypical anorexia nervosa presents a new avenue for determining the environmental component of complex chronic metabolic or metabolically-related diseases beyond the usual suspects of obesity and type 2 diabetes. In addition, the findings of this study provide a roadmap for investigating how the sustained application of our murine preconception exposure model in response to a wide range of metabolism disruptors can yield valuable insights into whether disparate exposures result in comparable offspring outcomes through analogous mechanisms or whether eukaryotic epigenomes possess the ability to transduce signals elicited by distinct exposures into functionally equivalent altered states.

## Acknowledgments

We extend our sincere gratitude to Daniel D. Davis for his invaluable contribution to the mouse dissections that underpinned this research.

## Data and code availability

The raw and processed data, as well as the necessary codes for reproducing analyses and figures, will be publicly distributed upon peer-reviewed publication.

## Author contribution

CDC and RCG conceived, designed and conducted the study, analyzed data and drafted the manuscript. SRA contributed to the mouse-related research.

